# Dual species interactions shield *Campylobacter* against multiple antibiotics

**DOI:** 10.64898/2026.02.27.708565

**Authors:** Sangeeta Banerji, Gudrun Holland, Michael Laue, Antje Flieger

**Author notes:** Corresponding author: Prof. Antje Flieger, Head Division of Enteropathogenic Bacteria and Legionella (FG11) and National Reference Centre for Salmonella and other Bacterial Enteric Pathogens, Burgstr. 37, 38855 Wernigerode, Germany.

## Abstract

**Background:** *Campylobacter* is a major food-borne pathogen causing diarrhea. In severe cases, treatment typically involves fluoroquinolones (FQ) and macrolides. However, high proportions of FQ resistance (FQR) have made macrolides the remaining first-line treatment option. As part of the *Campylobacter* surveillance program in Germany, we investigated multidrug resistance (MDR) in clinical *Campylobacter* samples collected 2010 to 2022.

**Methods:** A total of 6,980 samples was single-colony purified and analyzed for antimicrobial resistance phenotypes. Genome sequencing was performed for 2,912 samples, and antimicrobial resistance genes were mapped. Cultures showing MDR were further analyzed for contaminating DNA and studied using scanning electron microscopy (SEM).

**Findings:** We found that 453 (6%) *Campylobacter* samples were resistant to both FQ and macrolides. Two of the *C. jejuni* samples were resistant to antibiotics from ten different classes. Genome analysis revealed that these samples, despite being derived from single colonies, contained >10% *Enterococcus* DNA reads. SEM confirmed the presence of coccoid bacteria interspersed with spiral-shaped *Campylobacter*. Additional culture-based purification resulted in pure *C. jejuni* isolates that retained FQR but lost macrolide resistance. We further showed that presence of MDR Enterococcus spp. in the mixed samples protected *C. jejuni* from above-MIC (minimum inhibitory concentration) of several ribosome-targeting antimicrobials whereas pure *Campylobacter* were susceptible. These findings revealed that *Campylobacter* may gain antimicrobial resistance advantages through close interactions with *Enterococcus* spp.

**Interpretation:** MDR phenotypes in *Campylobacter* cultures may arise through close, dual-species interactions with other highly resistant intestinal bacteria, such as MDR *Enterococcus* spp. and may have practical implications for antibiotic treatment.

**Funding:** This study was funded by the German Ministry of Health dedicated to the National Reference Centre for *Salmonella* and other Bacterial Enteric Pathogens.

## Introduction

Campylobacteriosis is a major bacterial food-borne disease, with annual incidence rates of 64·9 and 19·5 per 100,000 population in Europe and the USA, respectively ^1,2^. Although the *Campylobacter* genus includes over 30 different species, *C. jejuni* and *C. coli* are primarily responsible for human infections. In healthy individuals, the infection is typically self-limiting and confined to the gastrointestinal tract. Symptoms include abdominal pain, (bloody) diarrhea, fever, nausea, and vomiting. In rare cases, extra-intestinal manifestations, such as bloodstream infections, can occur at varying prevalence and may still be underreported^3^. Post infection complications can include reactive arthritis and neurological disorders, such as Guillain-Barré-Syndrome and Miller-Fischer Syndrome ^4-6^.

Approximately one-third of the Campylobacteriosis cases in Germany are treated with antibiotics ^4^. Treatment depends on the antibiotic susceptibility of the strain and primarily involves macrolides or fluorochinolones (FQ) ^4,7^. However, more than 60% of *C. jejuni* and 70% of *C. coli* strains are resistant to FQ, leaving macrolides as the only first-line treatment option ^8^. As a result, FQR *Campylobacter* strains were included in the prior WHO high-priority list of pathogens requiring the development of effective new drugs ^9^. The recent emergence of *Campylobacter* strains carrying the transferable 23rRNA methylase gene *erm(B)*, which confers resistance to macrolides and lincosamides, has raised further concerns due to the increasing prevalence of MDR Campylobacter ^10^. Additionally, resistance genes of potential Gram-positive bacterial origin, such as optrA from *Enterococcus faecalis*, have been discovered in *Campylobacter*, suggesting transfer of DNA between these species ^11^.

Here we analyzed MDR in *Campylobacter* and further characterized two clinical FQR *Campylobacter* samples derived from patients with gastroenteritis, showing resistance to multiple other antibiotics, including macrolides. We found that these *Campylobacter* strains were closely associated with MDR *Enterococcus* strains, which, through dual-species interaction, provided protection against several ribosome-targeting antibiotics.

## Results

### Unusual MDR profiles of two clinical *C. jejuni* isolates with FQ and macrolide resistance

Due to high rates of resistance to first-line antibiotics, particularly FQ, ongoing surveillance of antimicrobial resistance, and especially MDR, is essential for the clinical management of Campylobacteriosis. Within the German *Campylobacter* surveillance program, we analysed 6,980 *C. jejuni* and *C. coli* isolates, predominantly obtained from stool samples of patients with gastroenteritis, collected between 2010 and 2022. Following single-colony purification, species identification was performed by PCR, and phenotypic susceptibility testing was performed using antibiotics representing 10 antimicrobial classes. Overall, 1,136 samples (16%) were fully susceptible, 4,128 (59%) were resistant to either FQ (3834/ 55%) or macrolides (294/ 4%), and 453 (6%) were resistant to both FQ and macrolides. Two *C. jejuni* samples, MC14-02991.B (male patient, 33 years) and MC20-00984.21 (male patient, 66 years), remarkably displayed resistance across the entire test panel. Among others, both cultures showed reduced susceptibility to ciprofloxacin, azithromycin, clindamycin, meropenem, tetracycline, and gentamicin (Tables 1 and 2). Notably, both exhibited meropenem resistance, a phenotype that has rarely been reported in *Campylobacter* ^12,13^.

**Table 1.**
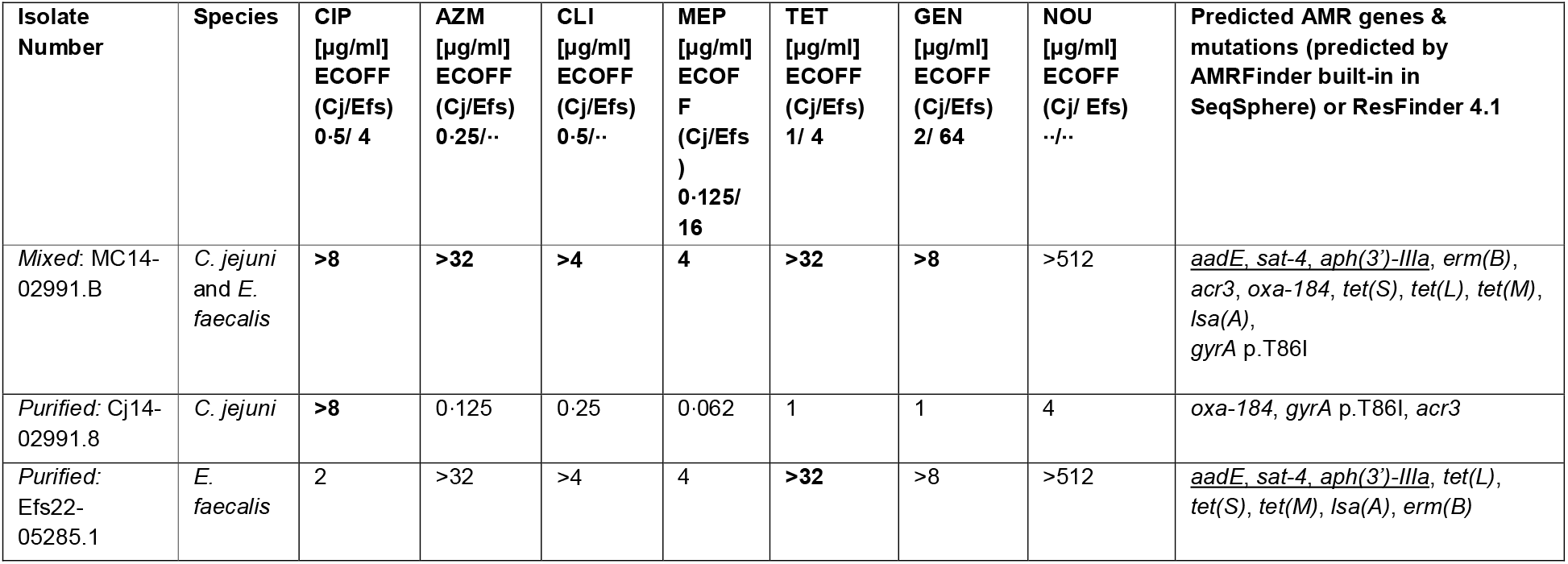
Resistance patterns of MC14-02991.B and derivatives Cj14-02991.8 and Efs22-05285.1. MIC values above the ECOFF for mixed samples and purified strains are marked bold and the respective isolates are categorized as non-wildtype for this antimicrobial. If a MIC value is below the ECOFF, the corresponding isolate is categorized as wildtype for this antimicrobial. Since there is no C. jejuni ECOFF for meropenem available from EUCAST, ertapenem ECOFF is given instead. Currently, no Enterococcus ECOFFS or breakpoints are available for azithromycin, clindamycin and nourseothricin (EUCAST) (see Material and Methods) and therefore the respective MIC values cannot be categorized into wildtype/ non-wildtype. The ASA cluster is underlined. See material and methods for full panel.

**Table 2.** Resistance pattern of MC20-00984.21 and derivatives Cj20-00984.4 and EFm22-05284.1. MIC values above the ECOFF for *C. jejuni* (Cj) or *E. faecium* (Efm) are marked bold and the respective isolates are categorized as non-wildtype for this antimicrobial. If a MIC value is below the ECOFF, the corresponding isolate is categorized as wildtype for this antimicrobial. Since there is no *C. jejuni* ECOFF for meropenem available from EUCAST, ertapenem ECOFF is given instead. If an ECOFF for *E. faecium* is missing the corresponding *E. faecalis* ECOFF is given instead. Currently, no *Enterococcus* ECOFFS or breakpoints are available for azithromycin, clindamycin and nourseothricin (EUCAST) (see Material and Methods) and therefore the respective MIC values cannot be categorized into wildtype/ non-wildtype. The ASA cluster is underlined. See material and methods for full panel.

| Isolate Number | Species | CIP<br>[µg/ml]<br>ECOFF<br>(Cj/<br>Efm)<br>0.5/ 8 | AZM<br>[µg/ml]<br>ECOFF<br>(Cj/ Efm)<br>0.25/ - | CLI<br>[µg/ml]<br>ECOFF<br>(Cj/ Efm)<br>0.5/ - | MEP<br>[µg/ml]<br>ECOFF<br>(Cj/ Efs)<br>0.125/ 16 | TET<br>[µg/ml]<br>ECOFF<br>(Cj/ Efm)<br>1/ 4 | GEN<br>[µg/ml]<br>ECOFF<br>(Cj/ Efm)<br>2/ 32 | NOU<br>[µg/ml]<br>ECOFF<br>(Cj/ Efm) -<br>/- | Predicted AMR genes &<br>mutations (predicted by<br>AMRFinder built-in in<br>SeqSphere) or ResFinder<br>4.1 |
| --- | --- | --- | --- | --- | --- | --- | --- | --- | --- |
| Mixed:<br>MC20-00984.21 | *C. jejuni*<br>and *E.*<br>*faecium*<br>mixed | >8 | >32 | >4 | >16 | >32 | >8 | >512 | <u>*aadE*</u>, *sat-4*, *aph(3')-IIIa*,<br>*tet(M)*, *tet(O)*, *oxa-193*,<br>*msr(C)*, *erm(B)*, *dfrG*,<br>*aac(6')-II*, *gyrA* p.T86I, *parC*<br>p.S80I, *pbp5_M485A*,<br>*50S_L22_A103V* |
| Purified:<br>Cj20-00984.4 | *C. jejuni* | >8 | 0.25 | 0.125 | 0.125 | >32 | 1 | >512 | <u>*aadE*</u>, *sat-4*, *aph(3')-IIIa*,<br>*oxa-193*, *tetO*, *gyrA* p.T86I,<br>*50S_L22_A103V* |
| Purified:<br>EFm22-05284.1 | *E. faecium* | >8 | >32 | >4 | >16 | >32 | >8 | >512 | <u>*aadE*</u>, *sat-4*, *aph(3')-IIIa*,<br>*msr(C)*, *erm(B)*, *dfrG*, *tet(M)*,<br>*aac(6')-II*, *parC* p.S80I,<br>*pbp5_M485A* |

### Detection of contaminating Enterococcal DNA by whole-genome sequencing

Next, 2,912 of the phenotyped strains were genome sequenced and, 8 (1·1 %) of the 747 macrolide-resistant *Campylobacter* samples showed the *ermB* gene known to confer resistance against macrolides, lincosamides, and streptogramin B (MLS_B_) including the two MDR samples resistant towards the whole test panel ^14^. The remainder displayed mutations in the 23S rRNA gene. Surprisingly, genome-based species determination identified contaminating *Enterococcus* DNA reads for both MDR samples, suggesting a contribution of the Enterococcal strains to the resistance phenotype. Indeed, reads from the first sample MC14-02991.B were attributed to both *C. jejuni* and *E. faecalis* and included resistance genes characteristic for both species, e.g. *tet(M), lsa(A)* typical for Gram-positive bacteria and *oxa-184* typical for *Campylobacter* (Table 1). Reads of the second sample MC20-00984.21 were attributed to both *C. jejuni* and *E. faecium* and included *tet(L), tet(M)* and *msr(C)* resistance genes characteristic for Gram-positive bacteria and *oxa-193* typical for *Campylobacter* (Table 2). Reads from the second sample MC20-00984.21 also revealed the *pbp5* gene with several mutations known to confer meropenem resistance (Table 2)^15^. After detecting *Enterococcus* DNA reads and resistance genes not previously described in *Campylobacter*, such as pbp5, we hypothesized that closely associated *Enterococcus* strains confer protection to *Campylobacter* during antibiotic exposure.

### Separating mixed-species cultures by serial subcultivation alters the Campylobacter resistance phenotype

Next, we sought to separate the *Campylobacter* strains from their associated *Enterococcus* strains. First, *Enterococcus* spp. from both mixed-species samples were readily recovered on blood agar under normoxic conditions, yielding *E. faecalis* strain Efs22-05285.1 from MC14-02991.B and *E. faecium* strain EFm22-05284.1 from MC20-00984.21 (Table 1 and 2). Second, *Campylobacter* selective agar (CCDA) under microaerophilic conditions enabled successful isolation of the *Campylobacter* strain Cj14-02991.8 (Table 1). This strain was not resistant to azithromycin or any other of the tested antibiotics except ciprofloxacin and ampicillin.

Isolation of the *Campylobacter* strain from mixed sample MC20-00984.21 under the same conditions was challenging, due to slow growth of *E. faecium* on CCDA. To obtain pure *C. jejuni* isolates from MC20-00984.21, 27 single colonies were randomly selected from CCDA, restreaked on CCDA, and incubated under microaerophilic conditions. Six of the 27 colonies remained mixed-species cultures, whereas the remaining 21 were confirmed as pure *C. jejuni* isolates and one clone was selected for further experiments (Cj20-00984.4) which showed no growth on blood agar under normoxic conditions. Notably, none of the 21 purified *C. jejuni* isolates derived from MC20-00984.21 was resistant to azithromycin, whereas, as expected, all six mixed-species colonies retained azithromycin resistance. This shift in resistance phenotype upon purification of both samples provided further indication that the unusual resistance profile was mediated by the closely associated *Enterococcus* strains (Table 1 and Table 2).

### Scanning electron microscopy (SEM) reveals close association of *C. jejuni* with *Enterococcus* in mixed samples

Since the *C. jejuni*/*Enterococcus* mixed-samples grew as homogenous white colonies on CCDA and on Brucella agar (FIG S1), we subsequently determined colony ultrastructure of the mixed-species samples MC14-02991.B and MC20-00984.21 compared to the purified strains by SEM. To this end, we studied single-colonies of the mixed samples. We observed that coccoid bacteria, representing *E. faecalis* or *E. faecium*, were closely attached to spiral shaped *C. jejuni* bacteria which showed a network of connecting flagella (FIG 1 A). In contrast, the purified strains contained one type of bacteria, either spiral shaped *Campylobacter* or coccoid Enterococci (Fig. 1B). The latter was also observed for reference strains, which were analyzed for comparison (FIG 1 C).

**FIG 1.**
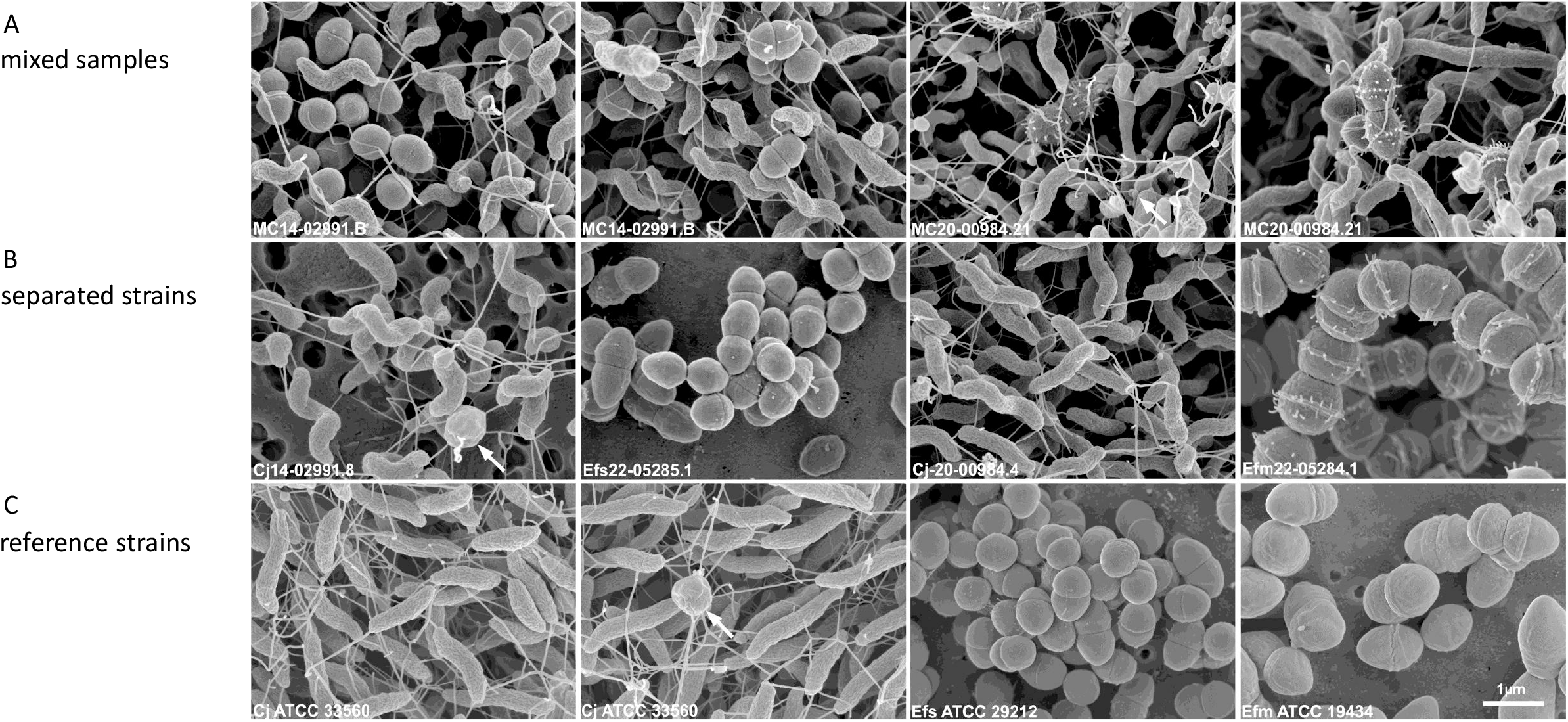
SEM demonstrates the association of *C. jejuni* with *Enterococcus* spp. in the same colony. FIG 1A: Images from two colonies each of mixed samples MC14-02991.B and MC20-00984.21 are shown. Bacteria were grown on *Brucella* agar or CCDA. For sample MC14-02991.B, coccoid bacteria were found in all investigated colonies of the mixed cultures. The cocci were either intermingled with *Campylobacter*, as shown here, or present in smaller groups. For MC20-00984.21, coccoid bacteria were found in some but not in all investigated colonies of the mixed samples. They were always intermingled with the spiral shaped *Campylobacter* and never appeared in groups. FIG 1B: Images from colonies of separated strains. From left to right: *C. jejuni* (Cj14-02991.8), *E. faecalis* (Efs22-05285.1), *C. jejuni* (Cj-20-00984.4) and *E. faecium* (Efm22-05284.1). Bacteria were grown on *Brucella* Agar or CCDA. On Brucella agar, *C. jejuni* may develop an unusual round to pleomorphic morphology (arrow) which can be the dominant form (see also Fig. 1C, Cj ATCC 33560). FIG 1C: Images from colonies of reference isolates. From left to right: *C. jejuni* (Cj ATCC 33560) grown on CCDA, *C. jejuni* (Cj ATCC 33560) grown on *Brucella* agar, E. faecalis (Efs ATCC 29212) grown on Brucella agar, *E. faecium* (Efm ATCC 19434) grown on CCDA.

### Associated Enterococcus strains account for elevated macrolide and meropenem MICs in mixed samples

Next, we compared the resistance phenotype of the purified isolates with the corresponding mixed-species cultures using broth microdilution assays. For each mixed sample, we selected one purified *C. jejuni* isolate (Cj14-02991.8 and Cj20-00984.4) together with the corresponding purified *Enterococcus* isolate (*E. faecalis* Efs22-05285.1 and *E. faecium* EFm22-05284.1). As expected, the mixed samples were resistant to all tested antimicrobials (Tables 1 and 2). Whereas, the purified *Campylobacter* strains were only resistant to ciprofloxacin (Cj14-02991.8) or ciprofloxacin, tetracycline and nourseothricin (Cj20-00984.4). Taken together, these data confirm that the elevated azithromycin and meropenem MICs observed in the *Campylobacter*/*Enterococcus* mixed cultures were attributable to the associated *E. faecalis* and *E. faecium*, respectively, whereas the corresponding purified *C. jejuni* cultures remained susceptible to these antibiotics.

### Whole genome sequence analysis reveals differential distribution of AMR genes typical for either *Campylobacter* or *Enterococcus*

The purified *C. jejuni* strains Cj14-02991.8, Cj20-00984.4 as well as the purified *E. faecalis* Efs22-05285.1 and *E. faecium* EFm22-05284.1 strains were subjected to whole-genome sequencing. Table 1 and Table 2 depict the detected antimicrobial resistance (AMR) genes and associated mutations. The analyses confirmed the differential distribution of AMR genes typical for either *Campylobacter* (e.g. *oxa-184, gyrA* p.T86I) or *Enterococcus* (*e*.*g. tet(M), lsa(A)*). Moreover, there were no significant contaminating Enterococcal DNA in the *Campylobacter* read samples. Further, the data interestingly indicated that strain Cj20-00984.4 and both Enterococcal strains shared the previously described *aadE-sat4-aphA-3* (ASA) cluster associated with aminoglycoside and streptothricin resistance, while strain Cj14-02991.8 did not contain it ^16,17^ (FIG S2). This finding underlines the presence of the ASA gene cluster both in *Campylobacter* and *Enterococcus*.

### Interactions with MDR *Enterococcus* enhances *Campylobacter* tolerance to several ribosome-targeting antibiotics

Next, we assessed whether intimate association with MDR *Enterococcus* strains enhances tolerance of *C. jejuni* to diverse antibiotics. Specifically, we examined whether *C. jejuni* within the mixed-cultures survived higher concentrations of different classes of antibiotics than pure cultures. To this end, we performed a microdilution and survival assay with the two mixed-species samples compared to the corresponding purified *Campylobacter* isolates (FIG 2). To distinguish these MICs, which suppressed cultivation of *C. jejuni*, from classical minimum inhibitory concentrations, we termed them isolation MICs. In the case of azithromycin, clindamycin, tetracycline (tetracycline: only for mixed strain MC14-02991.B), and gentamicin, we indeed observed a significantly enhanced survival of *C. jejuni* when associated with *Enterococcus* (FIG 3). However, no enhanced survival was observed for meropenem and nourseothricin (FIG 4). This shows that association with an MDR Enterococcus elevates resistance of *Campylobacter*, especially to various ribosome-targeting antibiotics. To assess whether this is a more general feature of *C. jejuni*/*E. faecium* interaction, we applied this assay on the type strain *C. jejuni* ATCC 33560 with and without co-incubation with MDR *E. faecium* strain EFm22-05284.1. We found that the type strain, too, showed elevated isolation MICs for the tested antibiotic azithromycin (see FIG. 5). Therefore, association with MDR Enterococci enhanced *Campylobacter* tolerance to antimicrobials.

**FIG 2.**
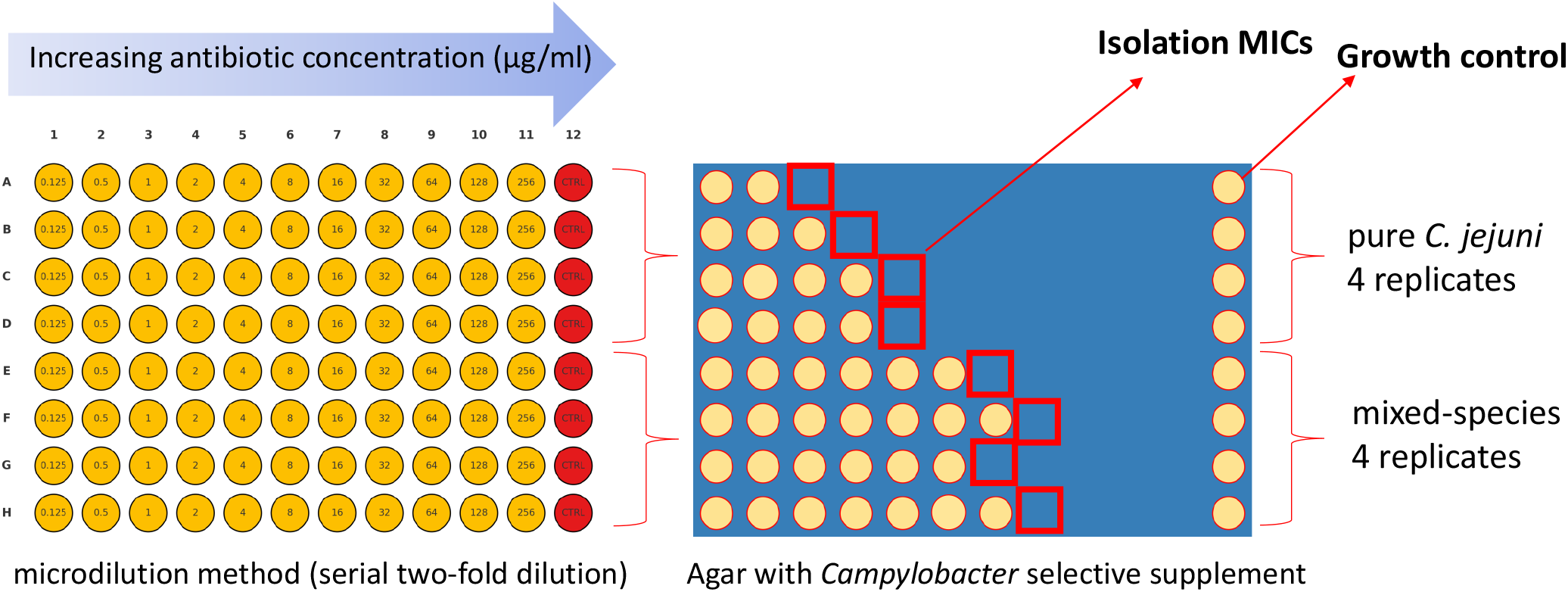
Workflow of microdilution assay for mixed-sample and pure *Campylobacter* MIC determination. First a microdilution assay was performed for every test antimicrobial for 24h incubation at 42°C (microaerophilic). For re-isolation of *C. jejuni*, 5 µl suspension from each well was transferred to Brucella agar containing *Campylobacter* selective supplement and incubated at 42°C for 48h under microaerophilic conditions. The agar did not allow growth of E. faecalis or sufficient growth of *E. faecium* within the given time of 48h. The isolation MIC is the lowest concentration of the antimicrobial, where no *C. jejuni* could be isolated from the wells, i.e. no growth was visible on the agar plate. The test was used for the following antimicrobials: azithromycin, clindamycin, tetracycline, gentamicin, meropenem and nourseothricin.

**FIG 3.**
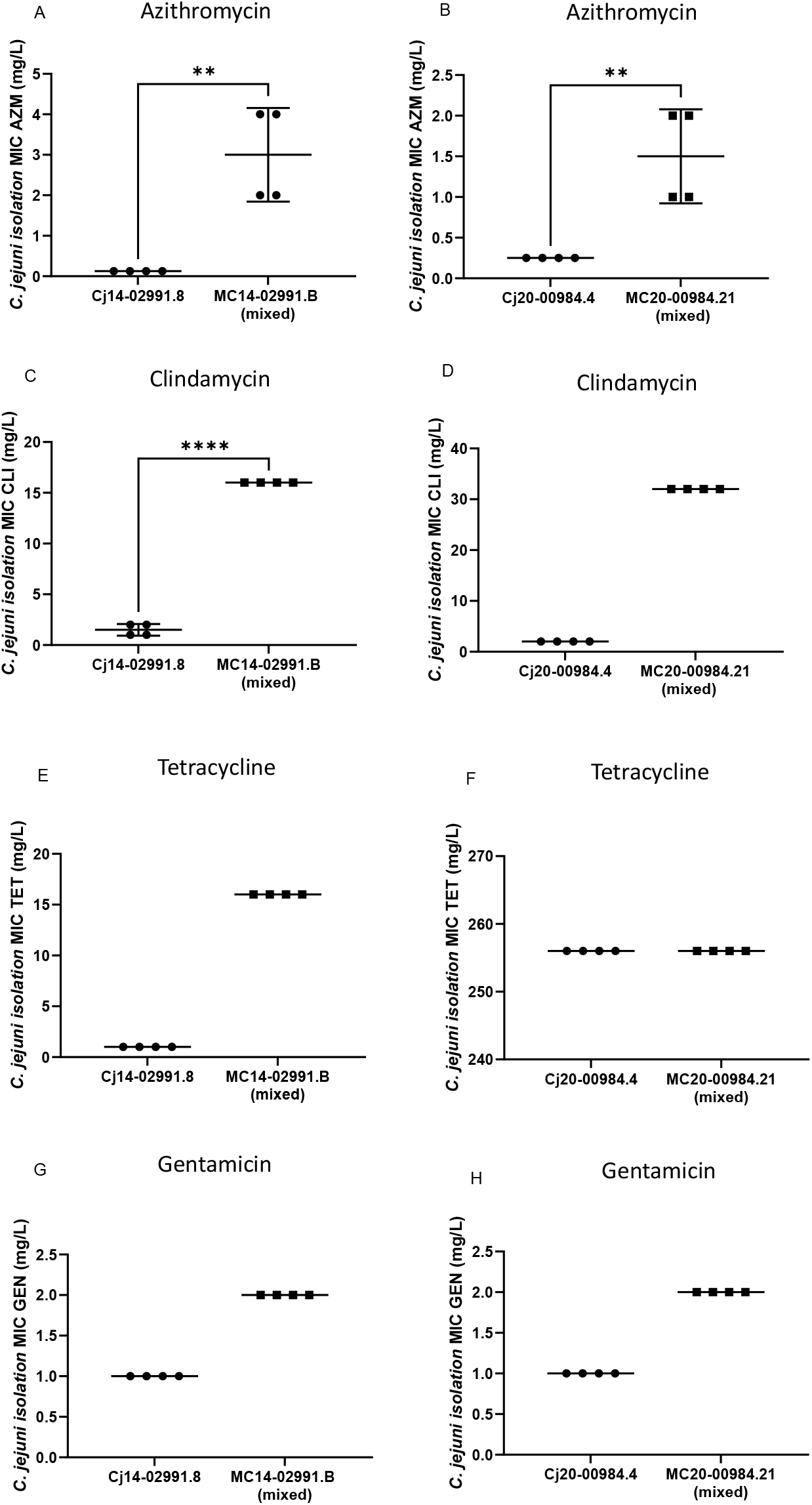
Interaction with *E. faecalis* or *E. faecium* protects *C. jejuni* against azithromycin, clindamycin, tetracycline, and gentamicin. One representative experiment of three experiments with four replicates each is shown. Error bars indicate standard deviation. asterisks indicate a significant difference (Student’s T-test, p<0·05). Please note for panels D, E, F, G, H: since all the values in each group were identical, it was not possible to compute a t test and no significance level could be calculated.

**FIG 4.**
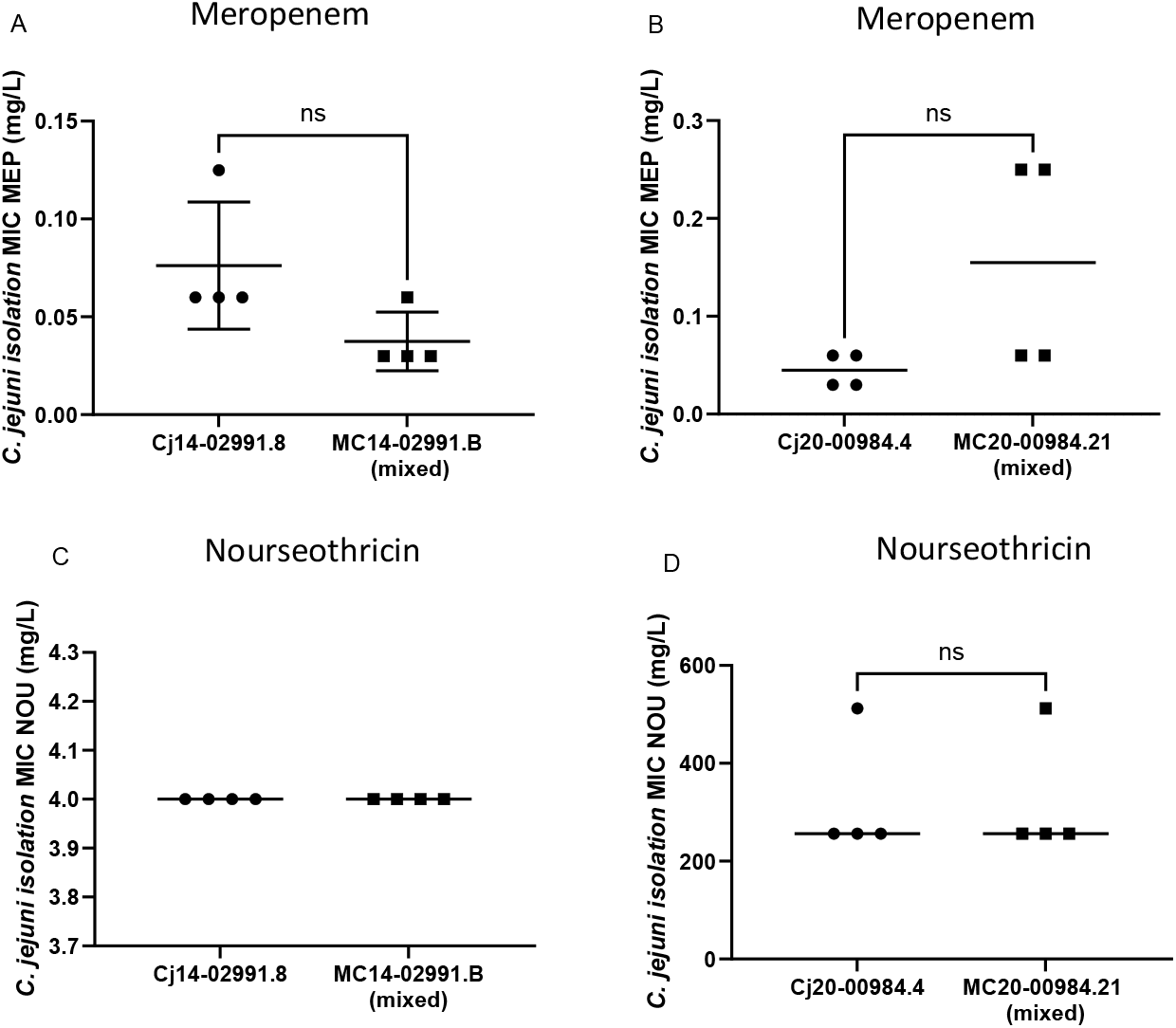
Interaction with *E. faecalis* or *E. faecium* does not enhance protection of *C. jejuni* towards meropenem or nourseothricin. One representative experiment of n=3 experiments with four replicates each is shown. Error bars indicate standard deviation. “ns” indicates a non-significant difference (Student’s T-test, p<0·05). Please note, since for panel C all the values in both groups were identical, it was not possible to compute a t test and no significance level could be calculated.

**FIG 5.**
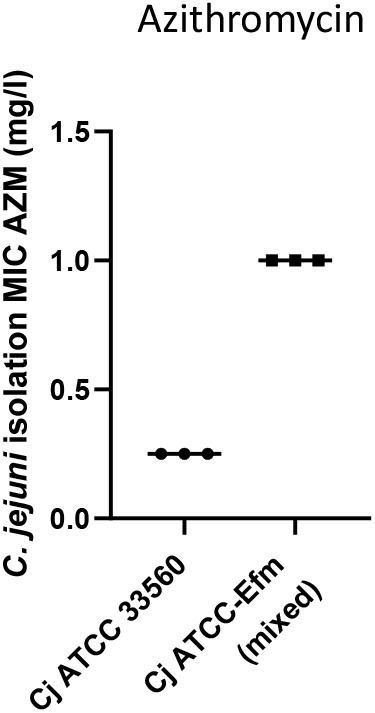
Interaction with *E. faecium* protects *C. jejuni* ATTC 33560 against azithromycin. One representative experiment of three experiments with three replicates each is shown. Error bars indicate standard deviation. Please note: since all the values in each group were identical, it was not possible to compute a t test and no significance level could be calculated.

### 0.7% of clinical *Campylobacter* genomes contain sequencing reads from contaminating species, suggesting that dual-species interactions occur in Campylobacter samples at a low but not negligible frequency

Last, we examined the prevalence of possible *Campylobacter* mixed-species samples in our genomic surveillance collection (2010-2022). 19 out of 2912 WGS sequences (0·7 %), including the two described above, indicated the presence of a major contaminating species (>10% foreign reads), predominantly Gram-positive bacteria of intestinal origin. Twelve of the 19 samples were aerobic species with 50% (6/12) being an *Enterobacter spp*. (Table 3). In seven of the 19 cases (41%), obligate anaerobic species were identified alongside *C. jejuni* or *C. coli*, such as Bacteroides *fragilis* (Table 3).

**Table 3.** Overview of *Campylobacter* mixed-species samples found by Mash (from 2912 sequenced samples from 2010-2022). AMR genes unusual for *Campylobacter* are highlighted in bold. AMR genes and mutations were predicted by AMRFinder built-in in SeqSphere or ResFinder 4.1.

| Isolate | *Campylobacter* species (identified by PCR + sequencing) | Other species (identified by sequencing) | AMR pattern (phenotypic) | Predicted AMR genes and mutations |
| --- | --- | --- | --- | --- |
| **Aerobic species** |  |  |  |  |
| MC14-02991.B | *C. jejuni* | *Enterococcus faecium* | AMK, AMP, CIP, CLI, CMP, ERY, GEN, KAN, NAL, OTE, STR, NAL, TET | <u>aadE</u>, <u>sat-4</u>, <u>aph(3')-IIIa</u>, erm(B), acr3, oxa-184, **tet(S)**, tet(L), tet(M), **Isa(A)**, gyrA p.T86I |
| 16-00111 | *C. jejuni* | *Enterobacter asburiae* | AMP, CIP, CLI, CMP, ERY, FOS, NAL, TET | **blaACT**, fosA, **oqxA/oqxb**, tetO |
| 16-00129 | *C. jejuni* | *Enterobacter hormaechei* | CIP, CLI, CMP, ERY, FOS, TET | **blaACT**, fosA, **oqxA/oqxb**, tetO |
| 16-00396 | *C. jejuni* | *Enterobacter asburiae* | AMP, CIP, CLI, ERY, FOS, TET | **blaACT**, fosA, **oqxA/oqxb**, tetO |
| 16-00573 | *C. jejuni* | *Enterobacter asburiae* | AMP, CLI, ERY, FOS, NAL, TET | **blaACT**, fosA, **oqxA/oqxb**, tetO |
| 16-00691 | *C. jejuni* | *Enterobacter asburiae* | CLI, CMP, ERY, FOS, TET | **blaACT**, fosA, **oqxA/oqxb**, tetO |
| 17-00289 | *C. coli* | *Enterobacter asburiae* | AMP, CIP, CLI, ERY, FOS, NAL, TET | **blaACT**, fosA, **oqxA/oqxb**, tetO |
| 19-02842 | *C. jejuni* | *Bacillus spizizenii* | AMP | blaOXA-603, **vmiR** |
| 19-07825 | *C. jejuni* | *Streptococcus parasanguinis* | AMP, CIP, FOS, KAN, NAL, TET, STR | aph3'(IIIa), blaOXA193, sat4, tetM, tetO, tetA(46) |
| MC20-00984.21 | *C. jejuni* | *Enterococcus. faecium* | AMK, AMP, CIP, CLI, CMP, ERY, FOS, GEN, KAN, MEP, NAL, TET, STR | <u>aadE</u>, <u>sat-4</u>, <u>aph(3')-IIIa</u>, tet(M), tet(O), oxa-193, **msr(C)**, erm(B), **dfrG**, **aac(6')-II**, gyrA p.T86I, **parC p.S80I**, **pbp5_M485A**, 50S_L22_A103V |
| 21-04765 | *C. coli* | *Morganella morganii* | AMK, AMP, CIP, CLI, ERY, FOS, GEN, MEM, NAL, STR, TC | <u>aph3'(IIIa)</u>, **blaDHA-1 (AmpC)**, **catA2**, tet(O) |
| 22-01023 | *C. jejuni* | *Staphylococcus epidermidis* | AMP, CIP, NAL, STR, TET | **fosB**, **dfrC** |
| **Anaerobic species** |  |  |  |  |
| 19-00899 | *C. jejuni* | *Bacteroides fragilis* (PCR confirmed) | FOS | blaOXA193 |
| 19-01673 | *C. jejuni* | *Alistipes shahii* (PCR negative) | AMP, CIP, NAL, TET | blaOXA193, **cfxA**, tetO |
| 19-02281 | *C. jejuni* | *Parabacteroides distasonis* (PCR confirmed) | AMP, CIP, NAL, TET | **blaOXA577**, **lnu/(An2)**, **erm(F)/mef(En2)**, tet(O), **tet(Q)** |
| 19-07765 | *C. jejuni* | *Phocaeicola vulgatus* (PCR confirmed) | susceptible | **blaOXA-61-like**, **cfxA** |
| 21-06223 | *C. coli* | *Parabacteroides distasonis*<br>(PCR negative) | FOS, GEN, STR | **aadE-Cc, CfxA, tet(Q)** |
| 22-00377 | *C. jejuni* | *Phocaeicola dorei* (PCR confirmed) | AMP, CIP, FOS, GEN, NAL, TET | **cfxA**, blaOXA-193, **erm(F)**, tet(O/32/O), **tet(Q)**, gyrA (p.T86I) |
| 22-04918 | *C. jejuni* | *Phocaeicola vulgatus* (PCR confirmed) | AMK, AMP, CIP, ERY, GEN. NAL, STR, TET | aph(3')-III, erm(B), sul2, **mef(A)**, 23S (g.2075A>G), gyrA_2 (p.T86I) |

In summary, this study identified close two-species interaction of two FQR *C. jejuni* and MDR *Enterococcus spp*. strains within the same colonies. This intimate association resulted in reduced susceptibility of *C. jejuni* towards multiple antimicrobials, especially ribosome-targeting antimicrobials. The identification of the mixed-species colonies was possible by means of WGS and SEM. WGS and laboratory data further indicated the co-occurrence of other species that may also have the potential for intimate association with *Campylobacter* and for mutualistic interactions influencing pathogen phenotypes

## Discussion

Isolation of bacterial strains from solid agar media based on single colonies was a vast achievement in infectious disease research and a basis for Koch’s postulates which link a pathogen and disease ^18^. It is concluded that all cells within such a colony originate from a single progenitor cell that undergoes repeated binary fission. All cells within a colony are assumed to be genetically identical; therefore, the colony represents a pure culture of microorganisms. Likewise, in clinical diagnostics, this principle is an important precondition for association of traits to a pathogen isolate, such as antibiotic resistance. Contradictory to the aforementioned, this study showed that *C. jejuni* and *Enterococcus* spp. can show very close interaction and accordingly are co-isolated from single colonies for repeated culture passages challenging Koch’s postulates.

The mixed-species cultures described here, comprising *C. jejuni* co-occurring with either *E. faecalis* or *E. faecium*, were recovered from stool samples of patients with gastroenteritis. Whole-genome sequencing and SEM indicated that spiral-shaped *C. jejuni* cells were intimately interspersed with coccoid *Enterococcus* cells, reminiscent of mixed-species microcolonies that can represent biofilm precursors^19^. SEM further suggested a meshwork of filamentous connections between neighbouring cells, consistent with flagella, which may contribute to the close association between the two species. In support of a central role of flagella in aggregate formation, Haddock et al. reported that *C. jejuni* forms microcolonies on human ileal tissue via flagella-mediated adhesion to mucus and to neighbouring bacteria, and that *flaA/flaB* mutants fail to form microcolonies or biofilms^20^. More broadly, *Campylobacter* flagella are increasingly recognized as multifunctional structures that contribute not only to motility but also to autoagglutination, adhesion and biofilm development. Collectively, the mixed-species aggregates observed here may represent remnants of intestinal biofilm-like communities and point to preferential interspecies interactions at this site.

Co-aggregation of *Campylobacter* with *Enterococcus* conferred a clear functional benefit, protecting *C. jejuni* from several ribosome-targeting antimicrobials. In mixed cultures, Enterococcus-associated resistance determinants, *ermB* (azithromycin/clindamycin), *tetS/tetM* (tetracycline) and aac(6⍰J)-Ii (gentamicin), were thus indirectly advantageous for *C. jejuni* survival, despite that these resistance mechanisms do not involve secreted drug-inactivating enzymes. Our data rather argue against acquisition of the genetic resistance markers by *C. jejuni* because the purified strains did not retain the resistance pattern of the mixed samples. Instead, we support involvement of a reversible, interaction-mediated phenotype. Further, azithromycin protection was reproducible in an independent strain background. Specifically, co-incubation of *C. jejuni* ATCC 33560 with the *E. faecium* isolate increased survival at above-MIC azithromycin concentrations, indicating that the protective effect is Enterococcus-driven (FIG 5). We therefore propose that close contact within microcolonies promotes regulatory changes in *C. jejuni*. For example, increased expression of the cmeABC efflux system enhances tetracycline resistance in *C. jejuni* and C. coli and could be triggered by diffusible danger signals such as peptidoglycan fragments released during lysis^21,22^. In line with this, co-incubation of *C. jejuni* with E. faecalis or E. faecium has been reported to alter Campylobacter protein abundance, potentially affecting cell–cell contact ^23^.

Resistance to fluoroquinolone antibiotics is frequently found in clinical *Campylobacter* strains leaving macrolides as the only other recommended first-line treatment option. In our study, we highlight the possibility that beneficial bacterial interaction can also provide protection against clinically important antimicrobials. Reduced antibiotic susceptibility of closely interacting bacteria, such as *Enterococcus*, could therefore influence resistance levels of *Campylobacter* which may have significance in the clinical setting.

When retrospectively analyzing the WGS data of 2912 samples collected between 2010 and 2022 in our strain collection, we found a contaminating species in 0.7 % of the *Campylobacter* samples. This included *Enterococcus faecium* and members of the *Enterobacter cloacae* complex. These are declared as WHO ESKAPE pathogens which can further have implications for antibiotic treatment in *Campylobacter* infections, which often is required in patients with underlying immune defect ^24^.

In conclusion, we found that gastroenteritis patients can be co-colonized with *C. jejuni* and MDR *Enterococcus* spp. resulting in an intimate association between the two species that confers protection to *C. jejuni* against diverse classes of antimicrobials, such as macrolides, lincosamides and others. Such interactions can provide the basis for unusual or extended MDR patterns, which so far have not been sufficiently explained ^29^.

## Methods

### Bacterial isolation and culture conditions

Single-colony *Campylobacter* spp. from patient stool samples were re-streaked for single colonies on *Brucella* agar (used until 2019) or *Campylobacter* blood-free agar (CCDA; from 2020) supplemented with C.A.T selective supplement (Oxoid; 8 mg/L cefoperazone, 4 mg/L teicoplanin, 10 mg/L amphotericin B). Strains were grown for 48–72 h at 42°C under microaerophilic conditions (5% O_2_, 11% CO_2_) as previously described ^25^.

Because CCDA usually inhibits *Enterococcus* spp., *Enterococcus* spp. (including ATCC29212 and ATCC19434) was cultivated on Brucella agar at 37°C under ambient air or at 42°C under microaerophilic conditions. In our set-up, *E. faecalis* did not grow on CCDA, whereas E. faecium showed slow and poor growth.

As *Campylobacter* spp. do not grow under ambient air, contamination was assessed by plating on blood agar and incubating aerobically (“blood agar test”). For colony morphology, bacteria were streaked on CCDA (to inhibit/reduce *Enterococcus* growth) or on *Brucella* agar (permits growth of *E. faecalis* and *E. faecium*) and incubated at 42°C under microaerophilic conditions for 48 h or 72 h.

Additional aerobic species (*Enterobacter cloacae, Bacillus subtilis, Streptococcus spp*., *Morganella morganii, Staphylococcus epidermidis*) were cultured on *Brucella* agar or blood agar at 37°C under ambient air or at 42°C under microaerophilic conditions. Additional anaerobic species (*Bacteroides fragilis, Phocaeicola vulgatus, Alistipes shahii, Parabacteroides distasonis*) were cultured on Brucella agar or blood agar at 37°C under hypoxic conditions (0.5% O_2_) or at 42°C under microaerophilic conditions.

### Identification of Campylobacter by means of PCR

*Campylobacter genus* and *C. jejuni/C. coli* species identification was performed by PCR using the primer sets described in US patent 2006/0051752 A1. *Enterococcus* genus and *E. faecalis/E. faecium* were identified by PCR using the species-specific assays recommended by the EURL-AR.

### *Campylobacter/Enterococcus* co-culture

*C. jejuni* ATTC 33560 was mixed 6:1 with *E. faecium* EFm22-05284.1 and grown over night at 42°C under microaerophilic conditions. The overnight culture was cryopreserved at −80°C and used for susceptibility testing and re-isolation of *C. jejuni*.

### Antimicrobial susceptibility testing

Biphasic liquid cultures (broth over an agar base) were incubated overnight without shaking at 42°C under microaerophilic conditions ^26^. Inocula were prepared from overnight liquid cultures in Brucella broth, adjusted in sterile saline to 0.5 McFarland, and diluted 1:1000 into wells of an in-house produced custom microtiter MIC panel. Panels were sealed and incubated for 24 h at 42°C in a humidified chamber under microaerophilic conditions (5% O_2_). Antibiotics (range, µg/mL) were: azithromycin (0.03–32), ciprofloxacin (0.063–64), nalidixic acid (4–32), tetracycline (0.5–8), ampicillin (1–16), gentamicin (0.5–8), kanamycin (2–32), amikacin (2–32), streptomycin (4–64), chloramphenicol (4–32), clindamycin (0.5–8), fosfomycin (8–128), meropenem (0.016–8) and nourseothricin (0.5–512). ECOFFs for *C. jejuni, E. faecalis* and E. faecium were applied where available. As EUCAST does not provide a meropenem ECOFF for Campylobacter spp., the ertapenem ECOFF (>0.125 mg/L) was used as a surrogate.

### Isolation of *Campylobacter* from broth microdilution assays

To re-isolate *C. jejuni* from broth microdilution wells, 5 µL from each well was spotted onto Brucella agar containing *Campylobacter* selective supplement (4 mg/L teicoplanin, 8 mg/L cefoperazone, 10 mg/L amphotericin B) and incubated for 48 h under hypoxic conditions (growth of *C. jejuni*; no growth of *E. faecium/E. faecalis* within 48 h).

### Short read genome sequencing and whole genome analysis

Short-read sequencing was performed in-house on an Illumina NextSeq 550 using the Nextera XT kit (up to 2×150 bp paired-end reads). Analyses were performed with Ridom SeqSphere+ v7.2.1 (Linux). Antibiotic resistance genes were identified using ResFinder 4.1 or AMRFinderPlus v3.11.2 integrated in SeqSphere+ v9.0.8 (Linux) ^27,28^. Mash (SeqSphere+) was used to identify contaminating reads; predicted genome size and average coverage were obtained from SeqSphere+. Mobile genetic elements and plasmids were assessed using MobileElementFinder v1.1.2 and MOB-suite v3.1.4 integrated in SeqSphere+ v9.0.8.

### Scanning electron microscopy (SEM)

Bacteria were grown on filters (PE Filter, 0.2µm, Pieper: PE02CP02500) placed on respective agar (MC14-02991.B, Cj14-02991.8, Efs22-05285.1 on Brucella agar and MC20-00984.21, Cj-20-00984.4, Efm22-05284.1 on CCDA) for four days under microaerophilic conditions. Filters were chemically fixed by floating on 1% formaldehyde and 2.5% glutaraldehyde (in 0.05 M HEPES buffer, pH 7.2) for 3h at room temperature. After dehydration with graded ethanol, filters were dried by critical point drying. and sputter-coated with 5 nm gold-palladium (Polaron Sputter Coating Unit E 5100,). Finally, the samples were examined with a field-emission SEM (Leo 1530 Gemini, Carl Zeiss Microscopy) at 3 kV using the in-lens secondary-electron detector.

## Acknowledgement

SB: Methodology, Validation, Formal analysis, Writing-Original Draft and Review, Visualization, GH: Investigation, Visualization (EM), ML: Resources, Methodology (EM), Writing-Review & Editing, AF: Conceptualization, Methodology, Resources, Writing-original Draft and Review & Editing, Supervision

This study was carried out as part of the German surveillance program for Campylobacter. We thank Christiane Schmidt for expert technical assistance. We thank the RKI unit for Nosocomial Infections and Antibiotic Resistance FG13 (Guido Werner, Jennifer Bender, and Yvonne Pfeifer) for the supply of Enterococcus reference strains. We further thank the laboratories who contributed to the surveillance program by submitting clinical Campylobacter strains.

## Data Sharing

This Whole Genome Shotgun project has been deposited at GenBank under the BioProject accession PRJNA1245490 and is publicly available. The Genome accessions are as follows: Efs22-05285.1: JBMRGQ000000000, MC14-02991.B: JBNQSB000000000, EFm22-05284.1: JBNPEJ000000000, MC20-00984.21: JBNQSA000000000, Cj14-02991.8: JBNJPZ000000000, and Cj20-00984.4: JBNJQA000000000.

