## Supplementary material for "Dual species interactions shield *Campylobacter* against multiple antibiotics"

Table of Contents

Figure S1: Page 1

Figure S2: Page 1

### Figures

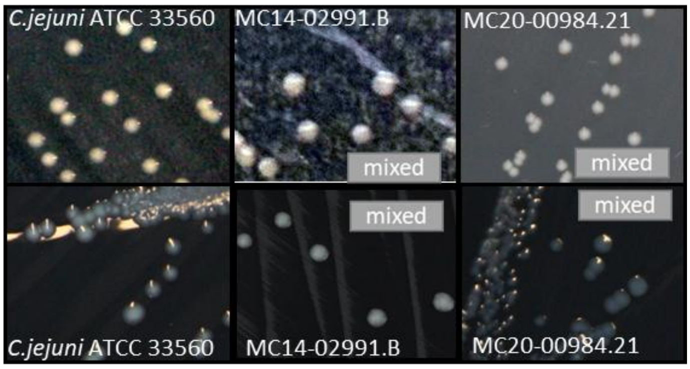


**FIG S1. Colony morphology of mixed-species samples resembles that of *C. jejuni* ATCC 33560 reference strain**

Mixed-species samples MC14-02991.B and MC20-00984.21 grown on *Brucella* agar (upper tiles) or CCDA (lower tiles) under microaerophilic conditions appear as homogenous colonies, which resemble in colour and shape that of *C. jejuni* reference strain ATCC 33560.

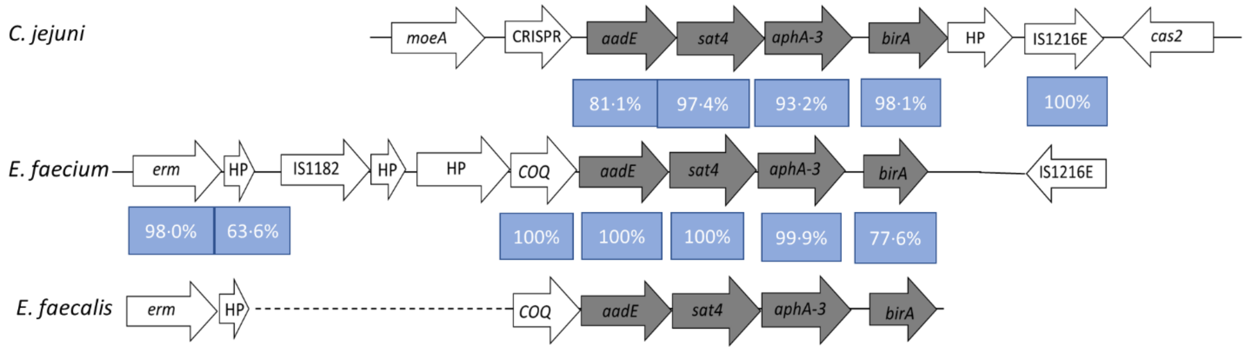


**FIG S2. Schematic overview of the genomic ASA cluster regions of *C. jejuni* Cj20-00984.4, *E. faecium* EFm22-05284.1 (reverse complement) and *E. faecalis* Efs22-05285.1.** The ASA Cluster region (marked in grey) is schematically visualized and the percent identity on nucleotide level between aligned homologous genes is shown in the blue fields. Solid lines represent the same contig and dashed lines represent detection of the genes on different contigs.
